# Potent inhibitors of toxic alpha-synuclein oligomers identified via cellular time-resolved FRET biosensor

**DOI:** 10.1101/2020.01.09.900845

**Authors:** Anthony R. Braun, Elly E. Liao, Mian Horvath, Malaney C. Young, Chih Hung Lo, Roland Brown, Michael D. Evans, Kelvin Luk, David D. Thomas, Jonathan N. Sachs

**Affiliations:** Department of Biomedical Engineering, University of Minnesota, Minneapolis, MN 55455; Department of Pathology and Laboratory Medicine, University of Pennsylvania, Philadelphia, PA 19104; Clinical and Translational Science Institute, University of Minnesota, Minneapolis, MN 55455; Department of Biochemistry, Molecular Biology and Biophysics, University of Minnesota, Minneapolis, MN 55455; Photonic Pharma LLC, Minneapolis, MN 55410

**Keywords:** Alpha-Synuclein, Toxic oligomers, Small-molecule inhibitors, Time-resolved FRET

## Abstract

Preventing or reversing the pathological misfolding and self-association of alpha-synuclein (aSyn) can rescue a broad spectrum of pathological cellular insults that manifest in Parkinson’s Disease (PD), Dementia with Lewy bodies (DLB), and other alpha-synucleinopathies. We have developed a high-throughput, FRET-based drug discovery platform that combines high-resolution protein structural detection in living cells with an array of functional and biophysical assays to identify novel lead compounds that protect SH-SY5Y cells from aSyn induced cytotoxicity as well as inhibiting seeded aSyn aggregation, even at nanomolar concentrations.

Our combination of cellular and cell-free assays allow us to distinguish between direct aSyn binding or indirect mechanisms of action (MOA). We focus on targeting oligomers with the requisite sensitivity to detect subtle protein structural changes that may lead to effective therapeutic discoveries for PD, DLB, and other alpha-synucleinopathies. Pilot high-throughput screens (HTS) using our aSyn cellular FRET biosensors has led to the discovery of the first nanomolar-affinity small molecules that disrupt toxic aSyn oligomers in cells and inhibit cell death. Primary neuron assays of aSyn pathology (e.g. phosphorylation of mouse aSyn PFF) show rescue of pathology for two of our tested compounds. Subsequent seeded thioflavin-t (ThioT) aSyn aggregation assays demonstrate these compounds deter or block aSyn fibril assembly. Other hit compounds identified in our HTS are known to modulate oxidative stress, autophagy, and ER stress, providing validation that our biosensor is sensitive to indirect MOA as well.

## INTRODUCTION

Misfolded and aggregated alpha-synuclein (aSyn) comprises the majority of the proteinaceous inclusions that are the pathological hallmark of Parkinson’s Disease (PD), Dementia with Lewy bodies (DLB), Multiple Systems Atrophy (MSA), and other alpha-synucleinopathies^*1–9*^. aSyn is a 140-amino acid, intrinsically disordered protein with a broad and diverse protein-protein interactome^*10–17*^. Multiple point mutations as well as gene duplication/triplication—resulting in the overexpression of wild-type (WT) aSyn—have been associated with Familial, inherited forms of PD and *DLB*^*18–24*^, however the majority of PD/DLB cases are sporadic, with no known direct genetic link to pathology^*25–27*^. The misfolding of aSyn into oligomeric and fibrillar has been shown to cause gain-of-function, toxic interactions with numerous protein targets (e.g. Tom20, 14-3-3, tau, ^*28*^ heat-shock proteins, amongst others)^*11, 13, 15, 29–41*^. Preventing or reversing the pathological misfolding and self-association of aSyn can rescue a broad spectrum of pathological cellular insults that manifest in PD, DLB, MSA, and other alpha-synucleinopathies.

There are competing yet synergistic hypothesis that describe the process of aSyn misfolding and subsequent pathology: the aSyn oligomer hypothesis and aSyn fibrillogenesis cascade. Lewy bodies are a shared, fibrillar, pathological hallmark of many alpha-synucleinopathies. These fibrillar inclusion have been posited to be contain neurotoxic species with a direct link to the prion-like, cell-to-cell spread of pathology throughout the central nervous system^*31, 42–47*^. Indeed, numerous rodent models have been developed using intramuscular or intra-striatal injection of sonicated pre-formed fibrils (PFF) and produce robust and temporally staged PD-like pathology^*44, 48–53*^. Although fibrils are present in the end-stage pathology of alpha-synucleinopathy, biochemical analysis of early stage transgenic animals suggest that the initial neuronal insult is associated with soluble aSyn oligomers, providing an additional pathological target for potential therapeutic intervention^*11, 15, 19, 42, 54–58*^.

Previous high-throughput screening (HTS) strategies targeting aSyn aggregation have been predominately focused on inhibiting or disrupting aSyn fibrils^*59–61*^. The aSyn fibrillogenesis cascade (**Fig. 1A**) is a complex process with a continuum of heterogenous aSyn assemblies that span intrinsically disordered monomers to rigid, β-sheet rich fibrils, with a range of on- and off-pathway oligomers with no-well defined secondary structure. Classically, the on-pathway oligomers are presumed to be early stage assemblies that nucleate and mature into β-sheet rich fibrils, whereas off-pathway oligomers do not feed into the fibrillogenesis cascade. Within the last few years, the importance of targeting oligomers as the pathological species has become forefront in the alpha-synucleinopathy research field^*9, 62–66*^ with increasing evidence that both on- and off-pathway oligomers are a heterogenous ensemble of oligomers comprised both toxic and non-toxic species.

**Fig. 1.**
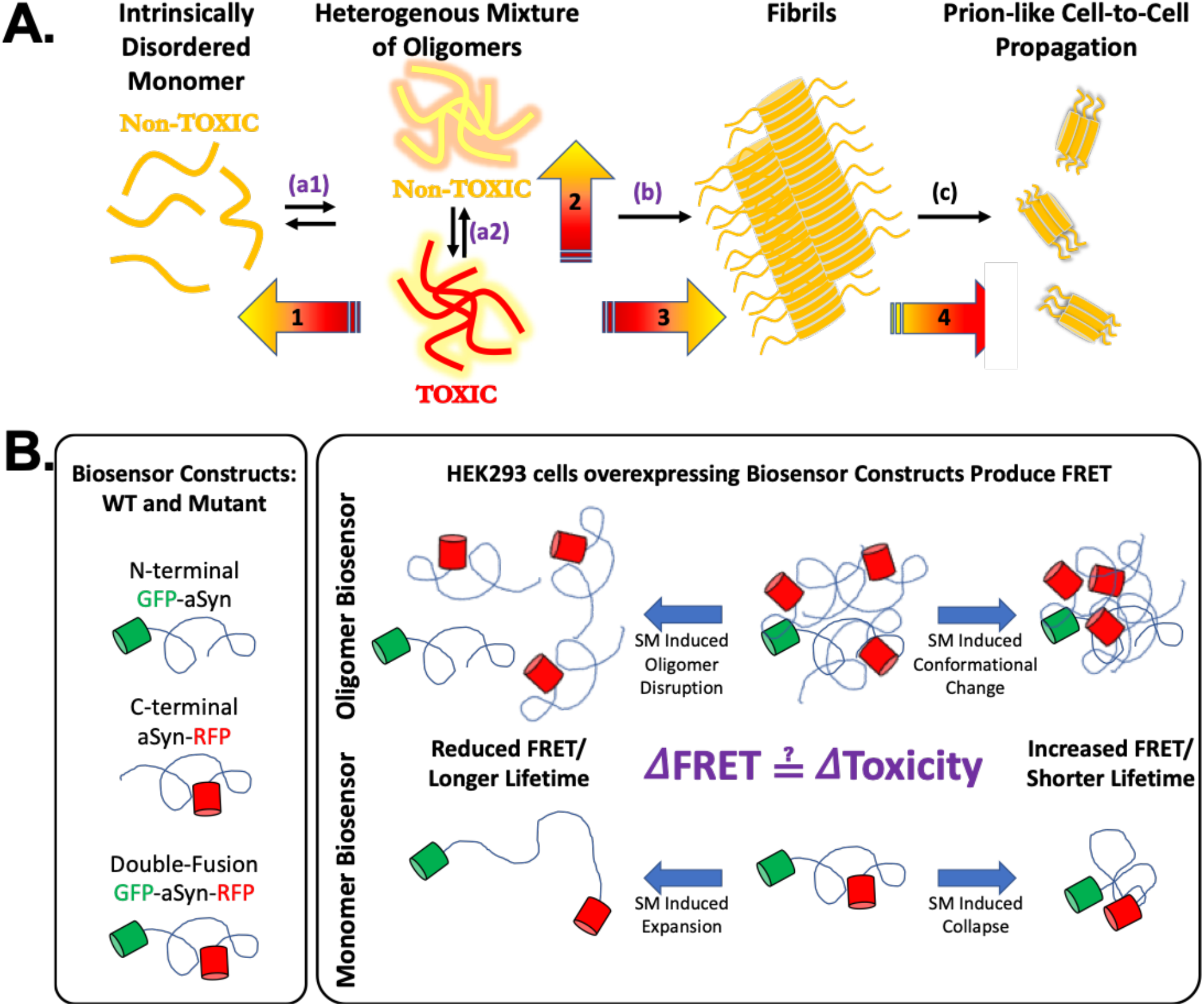
Monitoring spontaneous aSyn oligomer in live cell fluorescent lifetime biosensors. **(A)** The intrinsically disordered aSyn monomer is capable of misfolding into oligomers and fibrils, producing toxic assemblies that have been implicated in the pathology of disease. Oligomers comprise a heterogeneous mixture of metastable assemblies that have proven difficult to monitor with high precision and accuracy, so most current strategies target the irreversible formation of large assemblies (fibrils, arrow b). However, disruption of fibrils can induce toxicity due to elevated levels of toxic oligomers. Our cellular TR-FRET biosensors monitor reversible spontaneous oligomerization (a1) in addition to structural remodeling within the oligomer assembly (e.g. conversion between toxic and non-toxic oligomers, a2), while also monitoring downstream processes such as fibrillization (b) and seeded aggregation (c) with high sensitivity. The goal of this drug discovery pipeline is to identify compounds that modulate the toxicity of these aSyn assemblies (gradient arrows 1-4) and shift the equilibrium of this pathway toward non-toxic assemblies. **(B)** Intermolecular (oligomer) and intramolecular (monomer-conformation) cellular aSyn biosensors uses donor and acceptor labeled aSyn monomers that oligomerize when expressed in cells, producing FRET. Changes in FRET indicate dissociation and/or changes of oligomer structure (top) or a change in the ensemble of monomer conformations (bottom). These changes in FRET can be correlated to toxicity to link structure and function of aSyn oligomers.

The heterogenous nature of these aSyn oligomers have precluded more traditional biophysical techniques to characterize and resolve their distinct structure and properties. The McLean group developed a bimolecular fluorescence complementation (BiFC) assay that monitors aSyn oligomerization through the complementation and maturation of complementary truncated fluorescent proteins (*XFP*)^*67–69*^. This technique has provided direct *in vitro* and *in vivo* evidence of aSyn oligomers and their presence and enrichment in models of alpha synucleinopathies. However, the required maturation time and direct XFP-XFP interactions inherent in the BiFC assay limits its capacity to monitor spontaneous oligomer formation and subsequent oligomer disruption/remodeling by small molecules. This highlights the need for a novel platform to specifically target the spontaneous formation of aSyn oligomers without additional direct protein-protein interactions to facilitate HTS campaigns to identify small molecule inhibitors of toxic aSyn oligomer formation.

Fluorescent resonance energy transfer (FRET) provides a method to monitor protein-protein interactions without direct XFP-XFP contact. Numerous groups have employed FRET to study aSyn aggregation and monomer confirmation in both cell-free and cellular contexts^*28, 70–74*^ Recombinantly expressed, purified, and fluorescently labeled aSyn has been used in traditional FRET as well as single-molecule FRET experiments to elucidate oligomer and fibril formation under a wide range of conditions^*71, 72, 74–77*^. These systems have also been adopted for use in HTS campaigns to identify inhibitors of oligomer/fibril formation^*78*^. The simplicity and purity afforded by these recombinant protein systems are well suited for biophysical characterization of direct compound-to-aSyn mechanism of action (MOA). However, it has been well established that these synthetic, recombinant protein systems are far removed from the physiological context of aSyn misfolding and likely do not capture the essential protein-protein interaction environment (e.g. lacking the numerous chaperone proteins that interact with aSyn). Both single and double XFP-aSyn constructs have been used in cellular photo-bleaching FRET and FLIM studies to explore conformation of monomeric aSyn and punctate formation using a CFP/YFP FRET system. Cellular biosensors with these constructs have been developed by the Diamond group as biomarkers to probe brain lysate for fibrillar aSyn assemblies. These systems are focused on detection of pathological proteins in patient biofluids via seeded aggregation and do not focus on the early-stage spontaneous oligomerization of WT aSyn^*77, 79, 80*^.

Our drug-discovery platform includes a primary, cellular FRET based aSyn-oligomer biosensor coupled to an array of secondary/functional assays that are essential to determine a small-molecule’s potential mechanism of action (MOA). Our biosensor uses Fluorescence lifetime (FLT) based FRET to monitor spontaneously assembled, early-stage WT aSyn oligomers in live cells. Using FRET has added advantages over BiFC for monitoring aSyn oligomers due to the reporter molecule no longer requires direct physical interaction (as in BiFC) and having tunable R_0_ (Forster distance) for a given FRET XFP pair provides a range of distances with which we can optimize the biosensor. Furthermore, the use of FLT detection increases the precision of a FRET-based screening by a factor of 30 compared with conventional fluorescence intensity detection^*81*^ due to FLT’s intrinsic quality, dependent only upon the XFP and its local environment and not on the potentially highly variable XFP expression levels. This added sensitivity allows us to resolve minute structural changes within the ensemble of aSyn protein assemblies in the absence of seeded aggregation, providing a tool to monitor spontaneous oligomer formation and changes in the resulting oligomeric assemblies.

In this study we report results from a pilot HTS with of a 1280-compound library (LOPAC, Library of Pharmacologically Active Compounds) with our state of the art aSyn cellular FRET biosensors; through which *we discovered the first nanomolar-affinity small molecules that disrupt toxic aSyn oligomers in cells and inhibit cell death*. Subsequent ThioT seeded aSyn aggregation assays facilitated the identification of direct MOA. Other hit compounds that emerged from our pilot HTS have previously been shown to modulate oxidative stress, autophagy, and ER stress, providing validation our biosensor is sensitive to indirect MOA effects induced by aSyn. In addition, numerous hits have been previously identified to rescue aSyn induced toxicity providing further validation of our cellular FRET aSyn biosensor.

## RESULTS

### Biosensor Engineering

Our cellular aSyn biosensors monitor spontaneous oligomerization and the ensemble of monomeric conformations of aSyn. **Fig. 1B** is a schematic of our FRET biosensors. Their general design follows a similar development arc as our recently published tau biosensor platform^*82*^ with each aSyn construct being fused with either donor (N-terminal EGFP) an acceptor (C-terminal Tag-RFP), or both (EGFP-aSyn-TagRFP), all XFPs are monomerized forms.^*68*^ The decision to vary the termini for the XFPs was based on earlier BiFC assays developed by the McLean group^*69*^.

The core of our cellular bio-sensor system builds upon our established, state-of-the-art, HEK293/fluorescent lifetime plate reader (FLT-PR) platform designed to monitor specific protein interactions and structural changes *in living cells*.^*83–85*^ Fluorescence lifetime (FLT) detection increases the precision of FRET-based screening by a factor of 30 compared with conventional fluorescence intensity detection ^*81*^, and provides exquisite sensitivity to resolve minute structural changes within protein ensembles. The improved signal-to-noise of FLT detection is due to FLT’s being an inherent property of the XFP and not sensitive to fluctuations in steady state intensity. This sensitivity, along with our series of aSyn-XFP fusion constructs allow us to monitor inter-molecular aSyn interactions (oligomerization, **Fig. 1B**, *top*) as well as intra-molecular interactions (monomer conformation, **Fig. 1B**, *bottom*).

Initial biosensor characterization was performed via a titration of donor and acceptor DNA into HEK293 cells. Cells were transiently transfected and incubated for 48-hours, harvested, washed, and dispensed with 5-replicates for FLT characterization in our FLT-PR. **Fig. 2A** illustrates a robust donor only (1:0) FLT that undergoes increasing FLT attenuation with increased acceptor ratio through D:A of 1:12. There is a plateau in FLT reduction that correlates with overall donor-signal-to-noise. **Fig. 2B** presents the corresponding FRET for each system determined via Eq. 1, where there is significant increase in basal FRET up to the donor:acceptor ratio of 1:8.

**Fig. 2.**
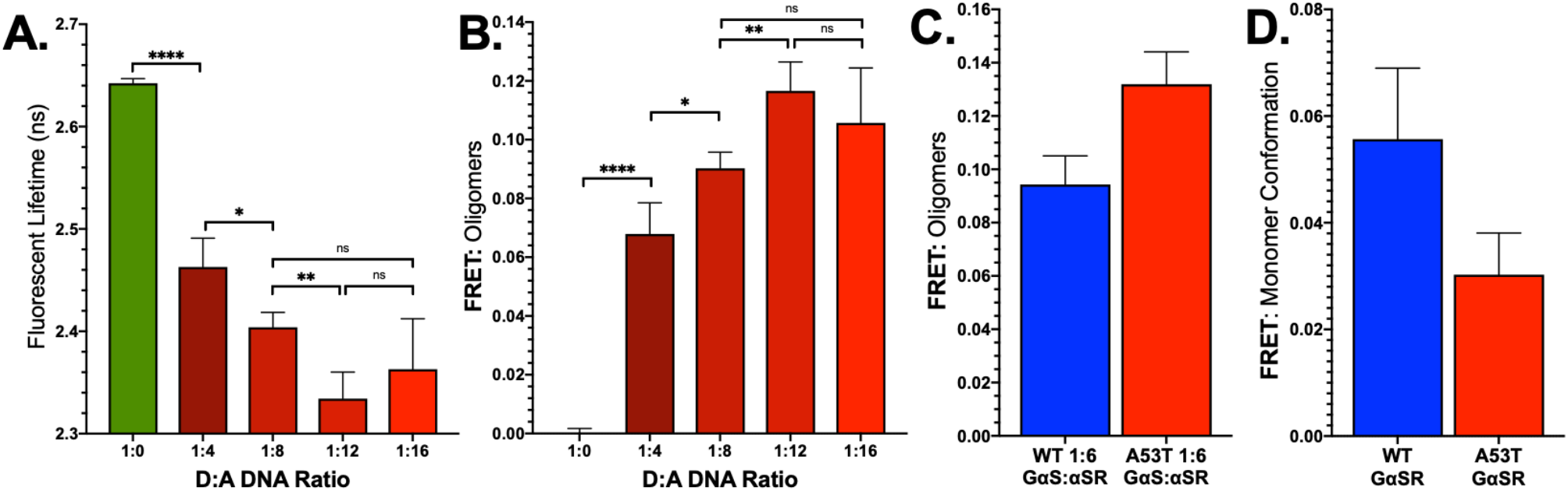
Characterization of aSyn cellular FRET biosensors. **(A)** Fluorescent lifetime measured in our FLT-PR for a range of donor and acceptor (D:A) ratios show significant reduction of FLT that saturates around D:A of 1:8. **(B)** FRET determined at each D:A ratio in panel A. **(C)** Comparison of WT and A53T aSyn oligomer biosensors show increased FRET for the aggregation prone A53T construct. **(D)** Interestingly, the double fusion intramolecular FRET biosensor that probes monomer conformation demonstrates a less compact monomer structure for the A53T mutant. *p < 0.05, ** p< 0.01, ****p<0.001, n.s. indicates not significant.

In addition to WT aSyn, we developed a set of mutant A53T aSyn biosensors to determine sensitivity of our biosensor system. The A53T familial mutant is known to increase aggregation propensity and toxicity. **Fig. 2C** and **Fig. 2D** compare WT and A53T FRET for both the oligomer and monomer biosensors respectively. As expected, the aggregation prone A53T mutant resulted in increased FRET relative to WT for the oligomer biosensor. In the monomer-conformation biosensor we observe reduced FRET, suggesting an extended conformation for A53T relative to WT.

Cellular biosensor expression was verified via fluorescent microscopy and Western blot analysis. **Fig. 3A** illustrates cells co-transfected with GFP-aSyn/aSyn-RFP, where the expression pattern is diffuse, devoid of puncta indicative of aggregated (fibrillar) aSyn. Western blot analysis of the cell-lysates from D:A ratios in **Fig. 2B** show distinct bands for both GFP-aSyn and aSyn-RFP (**Fig. 3B**). In addition to full-length aSyn fusion constructs, we also observe truncated aSyn species (more prominently in cells expressing the C-terminal RFP-aSyn fusion).

**Fig. 3.**
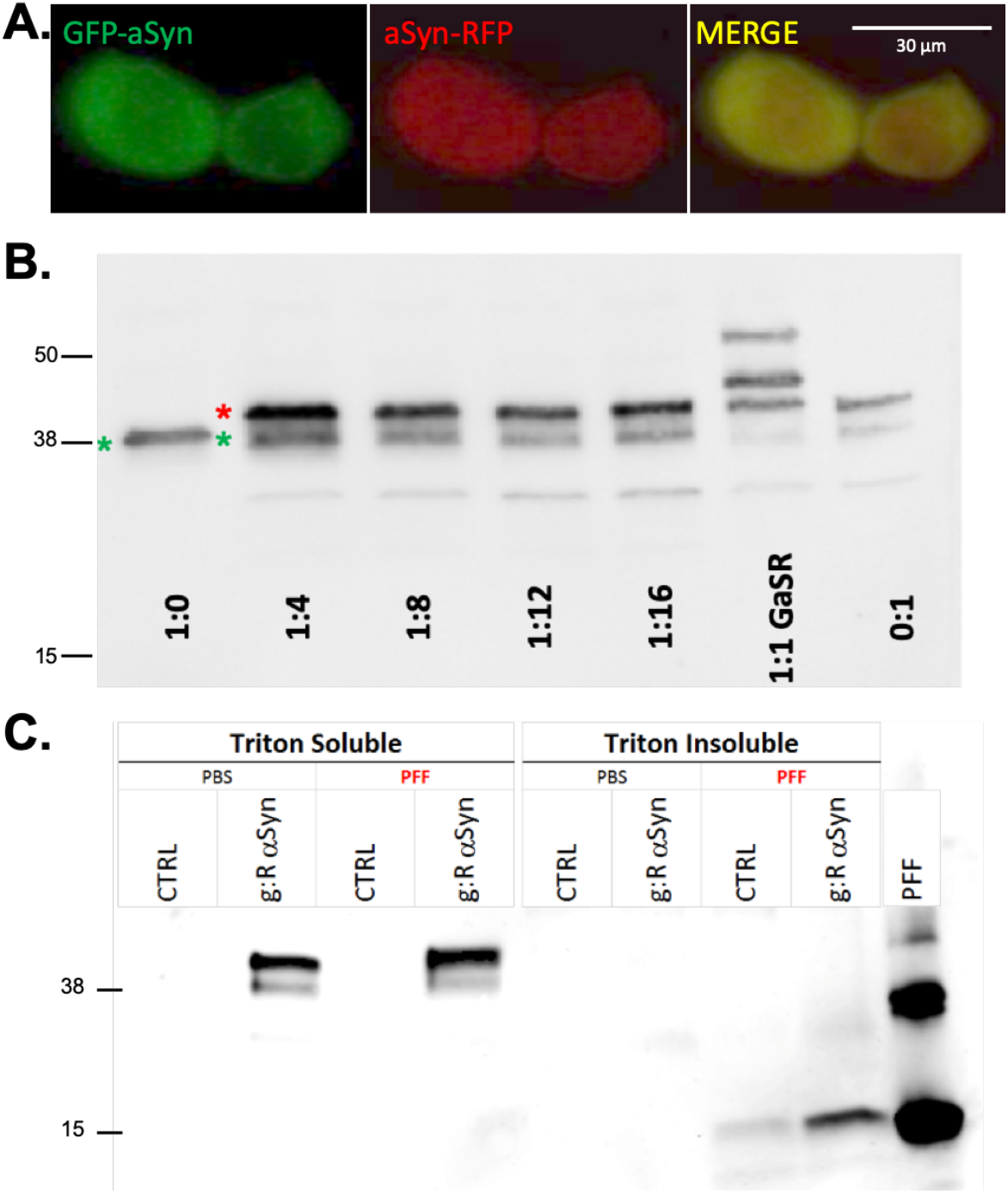
Characterization of aSyn cellular FRET biosensors. **(A)** Fluorescent live cell imaging of our cellular FRET biosensors co-expressing GFP-aSyn/aSyn-RFP constructs illustrate a diffuse expression profile (non-punctate). **(B)** Immunoblot for aSyn in WT aSyn oligomer (inter-molecular) biosensor. Cell lysates for each donor:acceptor ratio explored in Fig. 2A-B were analyzed via SDS-Page gel and probed for aSyn using the Syn101 antibody (BD labs). All samples were transfected with a constant total DNA (15uG DNA for 10cm plate). GFP-aSyn manifests as a strong band at ~40kD in the 1:0 lane. The C-terminal RFP-aSyn fusion runs slightly higher than the GFP-aSyn construct. Similarly, C-terminal truncation is observed for aSyn-RFP. Note: RFP truncation does not negatively affect our FRET assay as we only monitor the FLT for the donor (GFP-construct). **(C)** To explore whether our biosensor FRET stems from oligomers or fibrillary aSyn, both control and biosensor transfected HEK293 cells (g:R aSyn) were treated with freshly sonicated pre-formed fibrils (PFFs) or vehicle (PBS) for 24 hours and harvested with Triton-X soluble/insoluble extraction. All biosensor fusion constructs were extracted in the soluble fraction for both PBS and PFF conditions, indicating monomeric/oligomeric aSyn whereas PFF treated cells had insoluble aSyn, unlabeled aSyn.

Previous studies have shown that fibrillar alpha synuclein aggregates accumulate in the detergent insoluble fractions^*43, 86–88*^. Western blot analysis of Triton-X soluble and insoluble fractions of control HEK293 cells that were treated with PFF or vehicle (PBS) showed a modest increase in the amount of detergent insoluble aSyn, whereas HEK293 cells expressing the aSyn biosensor and treated with PFFs exhibited a threefold increase over controls in the amount of Triton-X insoluble aSyn (**Fig. 3C**). The experimental conditions for our aSyn biosensor HTS are more comparable to HEK293 cells that express aSyn treated with PBS. As seen in (**Fig. 3C**), there is minimal insoluble/fibrillar aSyn in the PBS treated samples. This provides evidence that our experiments are conducted in conditions that do not promote fibrillization of aSyn unless induced with aSyn PFF. Furthermore, the observed FRET from GFP-aSyn and aSyn-RFP is due to interaction of nonfibrillar aSyn assemblies (e.g. soluble aSyn oligomers).

### Pilot HTS with the LOPAC Library for the Oligomer and Monomer Conformation aSyn FRET Biosensors

Prior to each pilot HTS, FLT and emission spectrum were measured to verify signal level, basal FRET, coefficient of variance (CV) and similarity index. For each HTS, cells were dispensed into drug plates—1536-well (10μL/well)—and incubated with the compounds (10μM) or DMSO (1%) for 90min. FLT measurements were acquired with the FLT-PR. After deconvolving the instrument response function, a single exponential fit was used to determine the FLT for both the aSyn cellular FRET biosensor (τ_DA_) and the donor-only control biosensor (τ_D_). FRET efficiency was determined by Eq. 1. CV was determined from control plates (N=384 wells).

A major challenge to any fluorescent based HTS platform stems from potential fluorescent interference due to fluorescent compounds in the library. Fluorescent compounds (FC) were flagged using a spectral similarity index filter which evaluates the ratio of intensity from two bandwidths that span the donor emission spectrum (see ^*84*^ for more detail) and are excluded as potential false-positives^*84, 89–92*^. FLT for all compounds that passed the FC filter are presented in **Fig. 4** for both oligomer (**Fig. 4A**) and monomer-conformation (**Fig. 4B**) biosensor HTS. Each drug plate included 256-DMSO control wells as a control for the aSyn FRET biosensor signal quality over time. Both oligomer and monomer conformation HTS had a tight Gaussian distribution of compounds (**Supplemental Fig. 1** and **Fig. 4**). Compounds that changed FLT by more than 5 standard deviations (SD) were considered hits (**Fig. 4** red data points). The 5SD threshold is an arbitrary definition that was made to reduce the number of compounds that are investigated in subsequent secondary assays. For illustration a 3SD cutoff is indicated via green data points in **Fig. 4A** and **B**. In total 18-hits were identified with the oligomer (inter-molecular) and 8-hits with the monomer-conformation (intra-molecular) biosensor at the 5SD threshold (with 2-overlapping compounds for a total of 24-hits, subsequently labeled as A001 through A024).

**Fig. 4.**
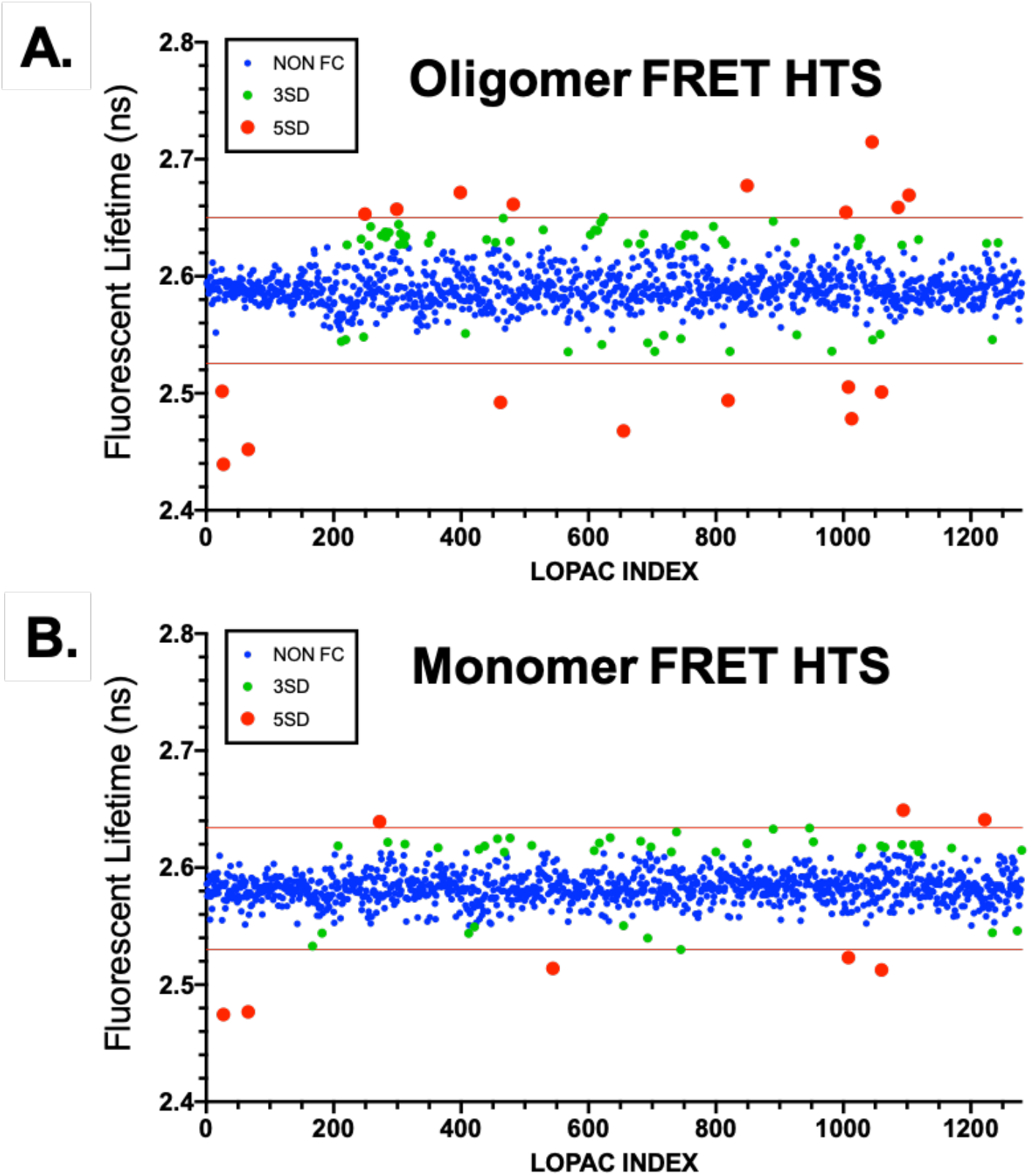
Pilot HTS for both Oligomer and Monomer cellular aSyn FRET biosensors. **(A and B)** Screens of the 1280 compound LOPAC library were performed in a single 1536-well plate for both the Oligomer (A) biosensor and Monomer conformation (B) biosensor. Pilot HTS provided 18 and 8 hits (oligomer and monomer biosensors, respectively) that changed FLT by > 5SD (red). For some systems a 3SD (red+green) threshold is sufficient if secondary assays can handle the increased throughput.

Our biosensor identified multiple compounds within the LOPAC library that were previously known to attenuate aSyn induced toxicity: AGK2 (a SIRT2 antagonist) and Bio (an autophagy inducer)^*93, 94*^. Additional hits span a range of known pharmacological functions that directly affect cellular pathways known to be disrupted by aSyn overexpression (e.g. modulating oxidative stress, reducing inflammation, inhibiting nitric oxide synthase, and upregulating autophagy). The identification of known effectors of aSyn pathology as well as compounds relevant to aSyn induced cellular dysfunction highlight the sensitivity and selectivity of our aSyn cellular FRET biosensor.

### FRET Dose Response of Hits with Wild-Type and A53T aSyn FRET Biosensors

Hits from the pilot HTS of the LOPAC library were selected and dispensed into 384-well plates to verify reproducible and specific FRET trends using the wild-type (WT) and A53T oligomer (inter-molecular) FRET biosensors. **Fig. 5** presents a subset of these dose response curves for Hit compounds A003, A008, A011, and A021; along with known control compound EGCG. For both WT and A53T biosensors the compounds displayed similar dose response signal. Concentrations above 50μM showed significant cytotoxicity for some compounds, prohibiting an accurate FRET IC_50_ determination. Of the hits tested, only A011 showed an increasing FLT trend, suggesting dissociation of oligomers (decreased FRET). All other hits (including control compound EGCG) displayed a reduced donor FLT (increased FRET), indicative of increased oligomer formation and/or remodeling into more compact oligomer conformation. Assay quality was determined by *Z*’ (Eq. 2) where a *Z*’ > 0.5 indicates good assay quality. Using EGCG as control compound and the hit A021 we calculated *Z*’ value of 0.76 and 0.66, respectively, indicating excellent assay quality.

**Fig. 5.**
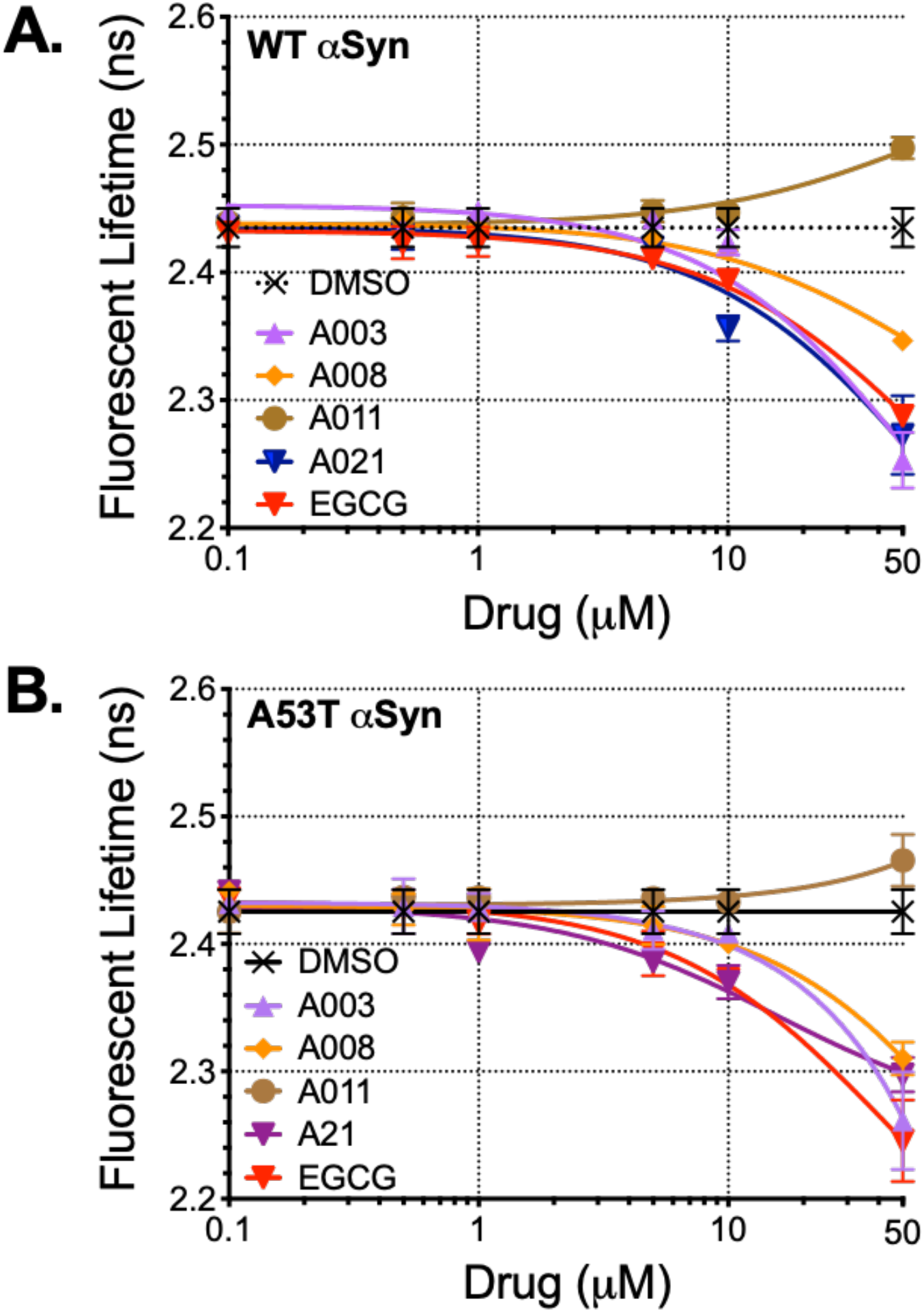
FRET dose response of pilot HTS hits. A subset of hits were evaluated using both WT **(A)** and A53T **(B)** oligomer biosensor to verify both increased FLT (disrupting aSyn-aSyn interactions) and decreased FLT (tighter aggregation or remodeled oligomers). Three hits that reduced FLT (A003, A008, A021) demonstrated similar response as control EGCG whereas hit A011 demonstrated mild recovery of FLT, suggesting dissociation.

### Secondary Assay: Total cellular viability

Hit compounds were evaluated for rescue of aSyn-induced cytotoxicity in using an SH-SY5Y neuroblastoma cell model of alpha-synucleinopathy. Overexpression of WT aSyn showed significant reduction in cell viability (46% total viability, **Fig. 6A** and **Supplemental Fig. 2**), whereas there is clear dose dependent rescue of toxicity for hits A003 and A021 with reduced effect for A008. Although hit A011 demonstrated reproducible and dosedependent increased in FLT (**Supplemental Fig. 2**), there was no observed effect on cytotoxicity, highlighting that a change if FLT does not necessarily correlate to changes in toxicity. A 7-point concentration response curve (CRC) for A003, A008, A021, and Anle138b (positive control compound, ^*95, 96*^) was performed to determine IC_50_ for each compound. Both A003 and A021 demonstrated potent inhibition of aSyn induced toxicity with an IC_50_ of **<u>78nM</u>** and **<u>65nM</u>**, respectively (**Fig. 6A**).

**Fig. 6.**
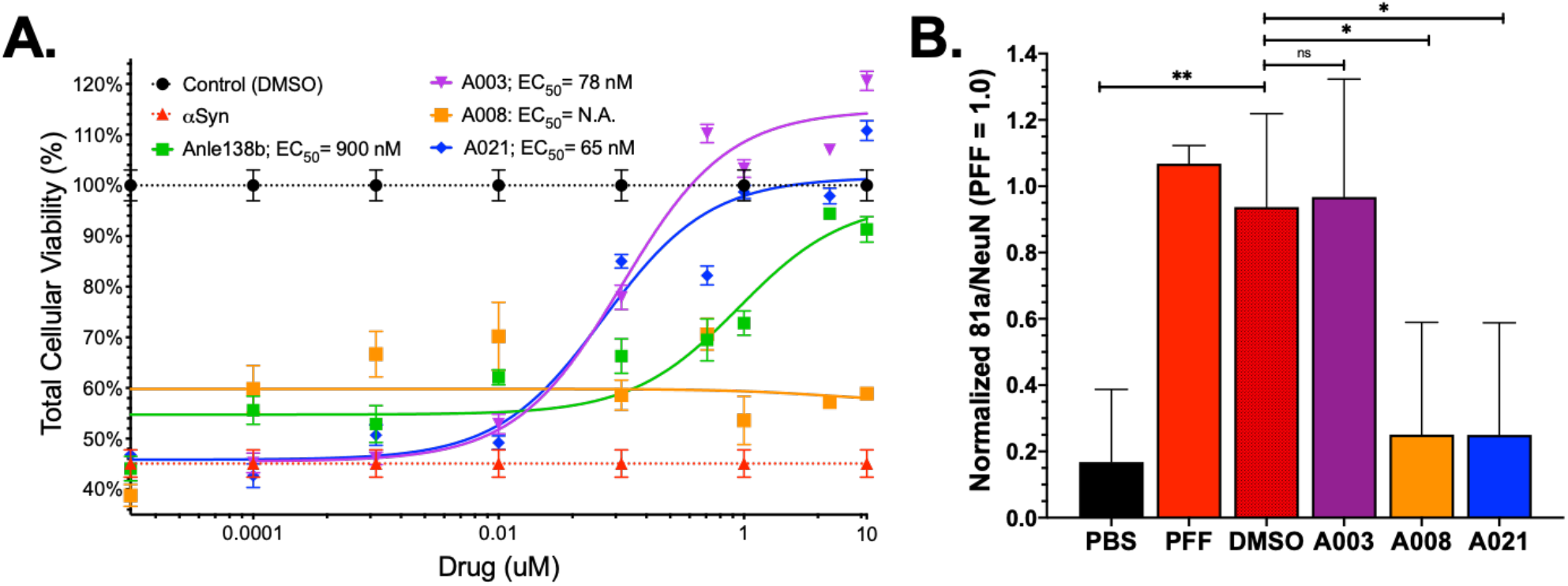
Cytotoxicity in SH-SY5Y cells and primary neuron pathology model demonstrate hit compound rescue of aSyn induce pathology. **(A)** Overexpression of aSyn in SH-SY5Y cells results in reduced cellular viability (~44%, red dashed line) as determined by CytoTox-Glo cytotoxicity assay. Treatment with control compound Anle138b as well as hit compounds A003 and A021 demonstrate a dose dependent rescue of toxicity with an EC_50_ of **900nM, 78nM**, and **65nM** for Anle138b, A003 and A021, respectively. Biosensor hit A008 did not rescue the SH-SY5Y cell toxicity. **(B)** Using an aSyn PFF induced pathology model in primary mouse neurons we evaluated the capacity of our hit compounds to rescue aSyn PFF associated pathology (phospho-S^129^ staining). Two of our hit compounds, reduced the aSyn pathology (A008 and A021). *p < 0.05, ** p< 0.01, n.s. indicates not significant.

### Secondary Assay: Inhibition of fibril growth

The aSyn fibrillogenesis cascade spans monomeric, oligomeric, and fibrils assemblies of aSyn. Thioflavin-T (ThT) aggregation assays have been robustly employed to monitor the kinetics and extent of fibril formation due to spontaneous and seeded aggregation^*59, 97–100*^. **Fig. 7** and **Supplemental Fig. 4** present results from a doseresponse seeded aggregation assay. Vehicle, control, and hit compounds were incubated with 15μM monomeric aSyn, doped with 5% freshly sonicated aSyn PFF in PBS under mild agitation (**Fig. 7A**). Seeded monomer samples underwent rapid elongation whereas monomer only samples did not spontaneous aggregate during the span of the experiment (**Supplemental Fig. 4B**). Titration of control and hit compounds were run in triplicate, representative traces of hit compounds are illustrated in **Fig. 7B** with a full summary of each titration detailed in **Supplemental Fig. 4C-H**. IC_50_ was determined for each compound by determining the max ThT intensity for each concentration (**Fig. 7C**). Both A003 and A021 demonstrated similar IC_50_ as control EGCG (931nM, 1.0μM, compared to 1.1μM for EGCG).

**Fig. 7.**
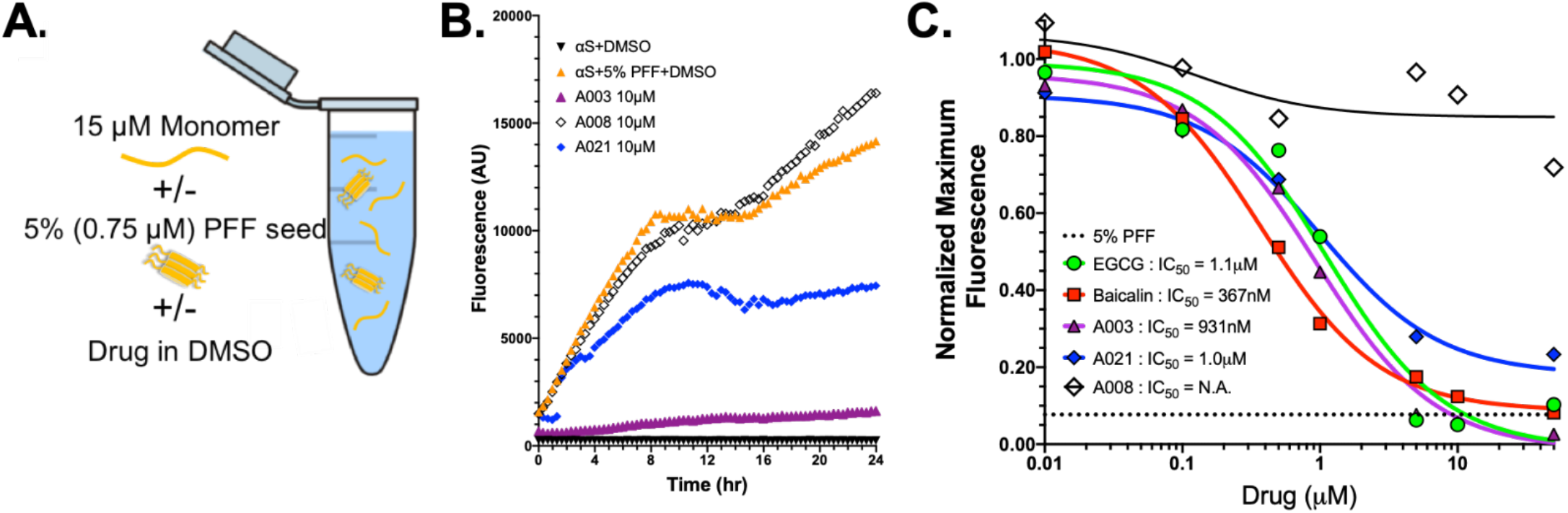
Seeded Thioflavin-T aggregation assay shows direct protein interaction for subset of hit compounds. **A)** Seeded ThT aggregation assay was performed using 15μM monomeric aSyn doped with 5% (0.75μM sonicated aSyn pre-formed fibril seeds, PFF) in a 384-well plate under slow agitation (300RPM) at 37C. **(B)** Control samples for the aggregation include PBS (black), aSyn monomer (brown), 5% aSyn PFF (red), and the 15μM monomer + 5% PFF mixture (orange). Dose response curves were made with a titration of compounds. Some hit compounds had little to no effect on aggregation, even at 50μM (e.g. A011). In contrast, A003 and A021 both show potent effects relative to Baicalin and EGCG controls. A008 does have some response but it lags behind other hit compounds. **(C)** Normalizing the initial peak in ThT data allows us to estimate an IC_50_ for each hit. Control compounds EGCG and baicalin show 1.1μM and 387nM potency whereas A003 and A021 have IC_50_ of 931nM and 1.0μM. Although there is a small trend with A008, the data cannot be fit to resolve an IC_50_.

### Secondary Assay: Primary Neuron Model of aSyn PFF pathology

Further analysis of hits A003, A008, and A021 in an aSyn PFF model also exhibited some protection against later stage aSyn pathology. As described previously, numerous *in vivo* and *in vitro* alpha-synucleinopathy mouse models have been developed using treatment with sonicated aSyn PFF to model pathology. PFF treatment of primary mouse hippocampal neurons results in neurotoxicity and increased in phosphoserine-129 (pS^129^) immunoreactivity as a marker for alpha-synucleinopathy pathology^*101, 102*^. These effects can be produced using either human-PFF or murine-PFF with the murine-PFF providing a stronger response. Primary neurons (7 DIV) were treated for 7-days with murine-PFF or PBS and hit compound or DMSO (compound vehicle) at 1μM. Neurons were fixed and stained for NeuN (neuronal nuclei), Map2 (neurite and neuronal morphology), and 81a (pS^129^) (see **Supplemental Fig. 3**). PFF treatment results in loss of neurons and a significant increase in pathology staining (e.g. normalized 81a/NeuN) that is unchanged with DMSO treatment (**Fig. 6B**). Concomitant treatment with 1μM A008 and A021 rescues the pathology phenotype however treatment with A003 has little to no effect. This compelling result indicates that some of the hits detected by our FRET biosensor may be able to offer protection against different stages of the aSyn cascade from oligomeric to fibrillar assemblies.

## DISCUSSION

Here we detail the development of a novel drug-discovery platform that monitors the misfolding and conformation of aSyn in cellular environments. Through leveraging the exquisite sensitivity of FLT based FRET, these cellular biosensors are sensitive to small structural changes with the ensemble of heterogenous spontaneously formed oligomers. Through engineering both WT and A53T familial mutant cellular FRET biosensor, we demonstrate that these biosensors are capable of resolving changes in oligomerization and conformation induced by a single point mutation. Furthermore, immunoblot analysis and fluorescent microscopy highlight that our cellular biosensor are expressed in a diffuse pattern and have little to no triton-x insoluble aSyn.

Using our FLT-PR technology, we performed a pilot HTS on the LOPAC library in a 1536-well format. Through coupling our primary HTS with cellular functional and biophysical secondary assays, we identified multiple novel hit compounds capable of rescuing aSyn induced cytotoxicity at nM level potency. Two of our hits, A003 and A021 demonstrated remarkable potency and will be prioritized for future hit-to-lead studies including medicinal chemistry and structure-activity-relationship analysis (SAR) as well as pre-clinical animal models of alpha-synucleinopathy.

Interestingly, amongst the multiple hits from the LOPAC HTS were compounds previously shown to rescue aSyn induced pathology in other models of alpha-synucleinopathy as well as modulate cellular pathways associated with aSyn overexpression (e.g. oxidative stress, autophagy, ER stress, etc.). This highlights our drug discovery platform is sensitive to small molecules capable of both direct and indirect MOA effects of aSyn spontaneous oligomerization and emphasizes the potential application of our technology.

Changes in FLT in the oligomerization biosensor can be indicative of multiple processes. An increase in the basal oligomer FLT could manifest from the dissociation of spontaneous formed oligomers or through structural rearrangement of oligomers that shift the donor/acceptor XFPs further apart. Similarly, a reduction in FLT could be due to increased oligomerization or structural rearrangement toward a more compact oligomer. For the monomer conformation biosensor the FLT changes are more clearly interpretable, with reduced FLT being more compact and increased FLT being more extended conformations. It is important to note that we cannot determine, a priori, whether an increase or decrease in FLT correlates to reduced toxicity, making it essential that the primary FRET HTS assay is coupled with secondary functional assays to elucidate changes in toxicity.

As an orthogonal secondary assay, we evaluated our hit compounds in a mouse primary neuron model of aSyn pathology. Here, mouse pre-formed fibrils (ms-PFF) treatment induces phosphorylation of serine-129, a signal indicative of aSyn pathology. This model is quite distinct from the spontaneous oligomerization of human aSyn that is monitored with our cellular FRET biosensors. It was rather surprising that, multiple hit compounds were successful at rescuing PFF induced pathology. This observation highlights the utility of our approach, specifically small molecules that perturb the conformation of monomeric or oligomeric aSyn can be capable of changing the phosphorylation state and pathology of exogenously treated fibrillar aSyn, providing greater potential therapy application for our drug discovery pipeline.

The coupling of our primary FRET HTS assay to secondary functional and biophysical assays provide a means to identify three novel classes of compounds (**Fig. 8**). Small molecules that modulate aSyn conformation or oligomeric state while 1) rescuing aSyn pathology through direct target engagement; 2) rescuing aSyn pathology through indirect MOA; and 3) altering aSyn conformation without changing pathology phenotype. Although two of these three classes of hits become potential leads for novel therapeutics, the third class of compounds can provide insight into domains and conformations of aSyn that are either not-relevant for aSyn pathology or lead to enhanced cellular dysfunction.

**Fig. 8.**
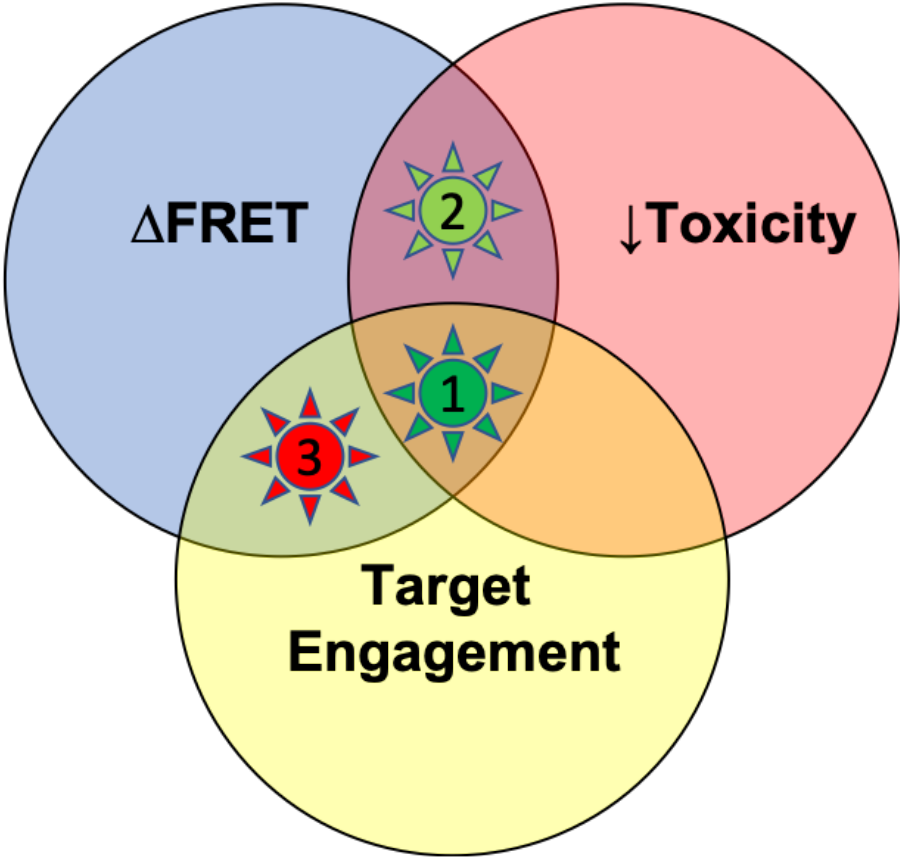
Novel Therapeutic Drug Discovery Pipeline for aSyn oligomerization is capable of elucidating multiple classes of novel compounds. Primary FLT FRET screen identifies compounds that induce a change in FRET, indicating changes in conformation or oligomerization state (blue). When coupled to secondary functional (cytotoxicity, red) or biophysical (target engagement, yellow), we are able to delineated three distinct classes of compounds that affect aSyn: 1) those that alter conformation, rescue toxicity, and directly engage aSyn; 2) those that alter conformation and rescue toxicity via indirect MOA; 3) those that change conformation but do not alter toxicity.

Our approach to developing this drug discovery pipeline has been consistent to previous efforts with other biosensor systems (e.g. tau and TNFR1, amongst others)^*84, 90–92, 103–105*^. At the core of any fluorescent assay is the potential of artifacts being introduced into the model system through the labeling of protein with synthetic fluorophores or fusion to large fluorescent XFPs. It would be naïve to ignore the potential concern that adding ~25kDa of XFP onto a 15kDa protein (as is the case for aSyn) can significantly alter the target protein’s native function. This is an ever-present challenge when developing fluorescent based assays and highlights the need for adequate controls and secondary, orthogonal assays that do not rely on the XFPs. As we evaluate cytotoxicity and MOA, we focus on unlabeled protein to minimize these potential artifacts.

Another concern with HTS campaigns is the physiological relevance of the cellular platform being deployed. The choice of HEK293 cells for our FRET HTS primary assay implementation is strongly rooted in the thorough vetting of this cell lines performance in multiple HTS campaigns that interrogated a wide range of protein targets ^*84, 90–92, 103–105*^. We acknowledge that not using neuronal cell lines in a primary HTS platform targeting neurodegeneration imparts a disconnect between the physiological and epigenetic link between cell type and disease environment. In this initial project, we work to mitigate this concern by implementing our secondary assays in a more physiological relevant SH-SY5Y or primary neuron model of pathology.

This does raise interesting questions that are central to many HTS strategies, what is the relevant standard that must be met to be provide relevant results that can be further developed toward lead compounds? If we are targeting a protein implicated in neurodegeneration, does the HTS platform need to be neuronal, human? What level of rigor is warranted or needed as a first-pass filter to identify compounds that directly interact with the target protein. These questions are not trivial to answer and are governed more so by available resources (reagents and funding) than actual scientific grounding. Nevertheless, we strive to improve our drug discovery pipeline by pushing toward more relevant model systems and reducing the assumptions and caveats inherent in our assays.

Lastly, in many HTS drug discovery campaigns, the focus on a hit compound MOA is as important if not more so than identifying the hit itself. Specificity and potency of potential therapeutics are directly related to whether or not the small molecule directly binds to the target of interest. Whether or not the small molecule behaves in a competitive or non-competitive manner for a particular interaction has strong implications on whether they may become a robust lead compound, destined for further medicinal chemistry optimization and clinical studies. These concepts are muddied in the aSyn protein-interaction space. The inherent promiscuity of the aSyn protein-interactome and its intrinsically disordered protein nature challenge our perceptions of what direct binding means for an IDP and how allosteric interactions can manifest and be modulated in an IDP.

In conclusion, our cellular FLT-FRET based aSyn biosensor found effective compounds that modulate oligomeric aSyn toxicity at nanomolar potencies and inhibited seeded aSyn fibrillization. The range of hits included molecules with potentially indirect MOA, where their known pharmacological targets directly affect known cellular dysfunctions associated with alpha-synucleinopathies. Lastly, although our cellular-FRET biosensor monitors the spontaneous oligomerization of non-fibrillar aSyn, we demonstrated that some of our hit compounds were also effective in decreasing, late stage, fibrillar aSyn pathology, highlighting the potential broad application of our drug discovery pipeline.

## METHODS

### Molecular biology

To generate the aSyn FRET constructs aSyn cDNA was cloned into a GFP-linker-RFP plasmid (linker contains 32 amino acids, GFP-32AA-RFP) that was previously characterized.^*106*^. QuikChange mutagenesis (Agilent Technologies, Santa Clara, CA) was performed to monomerize the GFP via A206K mutation and to produce the donor GFP-aSyn construct.^*107*^ Subsequent QuikChange mutagenesis was used to remove the 32 amino acid linker and stop codon after aSyn (creating GFP-aSyn-RFP construct) as well as to excise the GFP (creating unlabeled aSyn and aSyn-RFP constructs). The aSyn A53T mutation was introduced into all 4 plasmids. All biosensor plasmids constructs were sequenced for confirmation (ACGT, Wheeling, IL).

### Cell culture

HEK293 and SH-SY5Y cells (ATCC) were cultured in phenol red-free Dulbecco’s Modified Eagle Medium (DMEM, Gibco) supplemented with 2 mM L-Glutamine (Invitrogen), heat-inactivated 10% fetal bovine serum (FBS HI, Gibco), 100 U/ml penicillin and 100 μg/ml streptomycin (Gibco). Cell cultures were maintained in an incubator with 5% CO_2_ (Forma Series II Water Jacket CO_2_ Incubator, Thermo Scientific) at 37°C. The *inter*-molecular (oligomer) and *intra-molecular* (monomer-conformation) aSyn FRET biosensors were generated by transiently transfecting HEK293 cells using Lipofectamine 3000 (Invitrogen) with GFP-aSyn and aSyn-RFP (1:8 DNA plasmid concentration ratio) or GFP-aSyn-RFP plasmid, respectively The effectiveness of HEK293 cells transfected with FRET constructs as a HTS platform has been demonstrated in our previous work ^*84, 90–92, 103–105*^.

### LOPAC Library and Liquid Handling

The Library of Pharmacologically Active Compounds (LOPAC, Sigma-Aldrich) contains 1280 compounds that span marketed drugs, failed development candidates and naturally occurring compounds that have well-characterized activities with widely described biological effects. The library is originally formatted in 96-well mother plates and dispensed across four 384-well and/or one 1536-well flat, black-bottom polypropylene plates at 10 μM final concentration/well (50μL final volume for 384-well; 10μL for 1536-well) using an automated Echo acoustic liquid dispenser from Labcyte (Sunnyvale, CA, USA). DMSO (matching %v/v) was loaded as in-plate no-compound negative controls to make a total of 960 wells (384-well plates) or 256 wells (1536-well plate). The plates were sealed and stored at −20 °C until use.

Two days prior to screening, HEK293 cells were transfected using Lipofectamine 3000 with GFP-aSyn/aSyn-RFP (aSyn oligomer FRET biosensor) or GFP-aSyn-RFP (aSyn monomer conformation FRET biosensor) in 15 x 100 mm plates (6 x 10^6^ cells/plate). On each day of screening, the compound plates were equilibrated to room temperature (25 °C). The cells were harvested from the 100 mm plates by incubating with TrypLE (Invitrogen) for 2 min, washed three times in PBS by centrifugation at 200 x g and filtered using 70 μm cell strainers (BD Falcon). Cell viability prior to screen was assessed using a trypan blue assay, was >80%. Cells were diluted to 1.5E6 cells/ml. Expression of GFP-aSyn and the aSyn biosensors were confirmed by fluorescence microscopy prior to each screen (EvosFL-Auto brightfield microscope, Thermo Fisher Scientific, USA). After resuspension and dilution in PBS, the biosensor cells were constantly and gently stirred using a magnetic stir bar at room temperature, keeping the cells in suspension and evenly distributed to avoid clumping. During screening, cells (50 μl/well for 384-well; 10 μl/well for 1536-well) were dispensed by a Multidrop Combi liquid dispenser from Thermo (Pittsburg, PA, USA) into the assay plates containing the compounds and allowed to incubate at room temperature for 90 minutes before readings were taken by the fluorescence lifetime plate reader and spectral unmixing plate reader (Fluorescence Innovations, Inc) as described previously ^*108, 109*^. Reproducible FRET hits for retesting were purchased from Tocris (Minneapolis, MN, USA), Sigma (St. Louis, MO, USA), or Invitrogen (Carlsbad, CA, USA), depending on availability.

### HTS and fluorescence lifetime data analysis

As described previously ^*108, 109*^, time-resolved fluorescence waveforms for each well were fit with single-exponential decays using least-squares minimization global analysis software to give donor-acceptor lifetime (τ_DA_) and donor-only lifetime (τ_D_). FRET efficiency (*E*) was then calculated based on Equation 1.

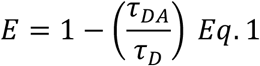

Assay quality was determined using control EGCG and hit compound A021 as positive controls and DMSO as a negative control and calculated based on Equation 2 ^*110*^,

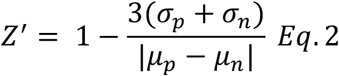

where σ_p_ and σ_n_ are the standard deviations (SD) of the observed τ_DA_ values, and μ_p_ and μ_n_ are the mean τ_DA_ values of the positive and negative controls.

Fluorescent compounds were flagged as potential false positives due to interference from compound fluorescence by a set of stringent fluorescent compound filters based on analysis of the spectral waveforms of each well from the LOPAC screen ^*108, 109*^. After removal of fluorescent compounds, a histogram of the FRET distribution from all compounds in the screen was plotted and fit to a Gaussian curve to obtain the mean (μ) and standard deviation (σ, SD). A hit was defined as a compound that decreased the FRET efficiency by more than five times the standard deviation (5σ) relative to the mean μ. (**Supplemental Fig. 1**)

### FRET dose-response assay

Compounds were tested in a FRET dose-response assay. The compound was dissolved to 10mM in DMSO which was serially diluted in 96-well mother plates. Each hit was re-screened for FRET dose response across eight concentrations (1 nM to 50 μM). Compounds (0.5 μl) were transferred from mother plates into assay plates using a Mosquito HV liquid handler (TTP Labtech Ltd, UK). The preparations for aSyn FRET biosensors were carried out as above.

### Cell cytotoxicity assay

Cell cytotoxicity was measured using the CytoTox-Glo (Promega Corporation) luminescence assay kit. SH-SY5Y human neuroblastoma cells were plated at a density of 5 x 10^6^ cells/plate in 100 mm plate (Corning) and transfected with unlabeled WT aSyn or equivalent vector-only control for 24 hours. The transfected cells were then plated at a density of 5000 cells/well in white solid 96-well plate (Corning) with a total volume of 100 μl, followed by treatment with hit compounds at three different concentrations (10 nM, 100nM 1μM), as well as DMSO-only controls, for another 48 hours. After incubation, 50 μl of CytoTox-Glo Cytotoxicity Assay Reagent was added to all wells followed by mixing by orbital shaking and incubation in the dark for 15 minutes at room temperature. Luminescence readings were measured using a Cytation3 Cell Imaging Multi-Mode Reader luminometer (BioTek). After the first read, 50 μl of Lysis Reagent was added, incubated for 15 minutes at room temperature, and a second read performed. Total viable cellular luminance was determined by the difference between the first and second luminescence signal and total cellular viability is determined by normalizing to the no-treatment samples.

### Western blot analysis

#### Biosensor Expression

To test the expression of aSyn FRET biosensors, HEK293 cells were plated in a six-well plate at a density of 1 x 10^6^ cells/well and transfected with GFP-aSyn/aSyn-RFP plasmids. Cells were lysed for 30 minutes on ice with radioimmunoprecipitation assay (RIPA) lysis buffer (Pierce RIPA buffer, Thermo Fisher Scientific) containing 1% protease inhibitor (Clontech, Mountain View, CA) and 1% phosphatase inhibitors (Millipore Sigma), and centrifuged at 15,000 g at 4 °C for 15 min. The total protein concentration of lysates was determined by bicinchoninic acid (BCA) assay (Pierce), and equal amounts of total protein (60 μg) were mixed with 4× Bio-Rad sample buffer and loaded onto 4%–15% Trisglycine sodium dodecyl sulfate–polyacrylamide gel electrophoresis (SDS-PAGE) gels (Bio-Rad, Hercules, CA). Proteins were transferred to supported nitrocellulose membrane and probed using Syn101 antibodies against aSyn (BD labs, San Jose, CA). Blots were imaged on using a ChemiDoc MP imager and analyzed using Image Lab (Bio-Rad, Hercules, CA).

#### Triton-X 100 Soluble/Insoluble Fractions

HEK cells were transiently transfected with empty vector or GFP-aSyn/aSyn-RFP as previously described. Cells were treated with either PBS or PFF (4μg/ml) 24 hours after transfection. After 24 hours of treatment, cells were harvested in cold lysis buffer (50 mM Tris/HCl pH 7.4, 150 mM NaCl, 5 mM EDTA pH 8.0) with protease and phosphatase inhibitors and 1% Triton-X 100 and incubated on ice for 30 min. The lysates were then centrifuged at 25,000xg for 60 min at 4°C. The supernatant was collected as the Triton-X 100 soluble fraction and the pellet was resuspended in lysis buffer with 2% SDS and sonicated for 10s and collected as the Triton-X 100 insoluble fraction. Both fractions were boiled and then a BCA protein assay was used to quantify the protein concentration of the samples. Approximately 12 μg of total protein were loaded onto a 4-20% mini Protean TGX gel (Bio-Rad, Hercules, CA) and blots were evaluated as described previously

#### Protein purification

Full-length wild-type aSyn protein was purified from BL21(DE3) *E. coli* using previously published protocols^*111*^. Protein production was induced with 320 μM IPTG for 8 hours. Cells were pelleted, lysed, and cut twice with Ammonium Sulfate. Subsequent purification was performed using an AKTA FPLC for both anion exchange and SEC purification. SEC fractions were assayed for purity, pooled, and concentrated in 3kd MWCO AmiconUltra spin columns. Protein concentration was determined via BCA, monomeric protein was then aliquoted at 5mg/mL (350 μM), and flash frozen and stored at −80 until use.

#### Pre-Formed Fibrils (PFF)

Bulk pre-formed fibrils (PFF) were following previously established protocols^*43*^. Briefly, aliquots of recombinant monomeric aSyn were rapidly thawed and spun at 25,000xg for 30min at 4C. For bulk PFF production approximately 500μL of 5mg/ml monomeric protein was added to a 2-ml Eppendorf tube and placed on a thermomixer-C (Eppendorf, USA) shaker and incubated at 37°C and 1000RPM for up to 144 hours. Every 6 hours, 5μl aliquots were removed and assayed for Thioflavin-T fluorescence until a plateau in aggregation was observed and the tube was opaque. 20uL aliquots of 5mg/mL PFF were then flash frozen and stored at −80 °C.

#### Thioflavin-T (ThT) assay

Thioflavin-T (ThT, Sigma, product no. T3516) was dissolved in PBS buffer and was filtered through a 0.2 μm syringe filter to make a stock solution of 2.5 mM. Working ThT concentration was 20 μM. Pre-formed fibril (PFF) seeds were produced via constant agitation of 1000 rpm at 37°C (Thermomixer-C, Eppendorf, Hauppauge, NY) of 500 μL of 350 μM aSyn in PBS in a 2 mL tube for 5 days. PFFs were aliquoted and flash frozen and stored at −80°C until further use. PFF seeds were rapidly thawed, diluted to 35 μM in PBS and sonicated 120 pulses at 50% using a probe-tip sonicator following established protocols^*43*^. Master mix samples for the ThT assay were prepared—PBS, 15 μM monomeric aSyn +/- 5% PFF (0.75 μM), 15 μM PFF or 0.75 μM PFF—in 20 μM ThT and plated in triplicate as 50 μL in 384-well flat, black-bottom polypropylene plates (PN 781209, Greiner Bio-One) and sealed with fluorescently transparent film. All empty wells were filled with PBS for a thermal sink and to help prevent evaporation. Assay plates were prepared as pre-cellular FRET dose response above. Dose response was done across 8-concentration (10nM to 50 μM). The assay was run at 37°C with mild shaking (200 rpm) in a Cytation3 plate reader. The ThT fluorescence was monitored by top-read with excitation filter of 440 nm and emission filter of 480 nm. Readings were acquired every 20 minutes for a minimum of 24 hours.

IC_50_ values were determined by first normalizing all ThT curves to the maximal signal at plateau for the aSyn+5%PFF+DMSO samples. The normalized fluorescence at each concentration was fit to the Hill equation to resolve an effective IC_50_ of seeded aggregation inhibition.

#### Primary hippocampal neuron cultures and fibril transduction

Primary neuronal cultures were prepared from CD1 embryos on E16-18, as previously described^*43*^. Tissue culture plates and coverslips were coated with Poly-d-lysine (Sigma) before addition of cells. Neurons were plated in 96-well plates (60,000 cells/cm^2^). Cultures were maintained in Neurobasal medium supplemented with B27 and Glutamax (all from Invitrogen).

PFF treatment was performed at 7 days in vitro (DIV). Briefly, stock aSyn PFFs were diluted in sterile PBS (without Ca^2+^/Mg^2+^; Corning) and sonicated with a bath sonicator (Bioruptor Plus, Diagenode) for 10 cycles at high power (30s on, 30s off, at 10°C). Sonicated PFFs were then further diluted to the indicated final concentration in neuronal media and added to cultures. PFF concentrations are expressed as the total equivalent aSyn monomer content in the preparation. Cultures were treated with 200 nM PFFs unless otherwise noted. Compounds were initially diluted in neuronal media before being added to the PFF neuronal media solution at a final concentration of 1-10 μM.

#### Immunocytochemistry and antibodies

Cultured neurons were fixed by replacing media with warm paraformaldehyde (4% in PBS containing 4% sucrose) for 15 min at room temperature (RT). Fixed neurons were washed three times with PBS and then blocked (3% bovine serum albumin, 3% fetal bovine serum in PBS) for 1h at RT. Cells were then incubated in primary antibodies diluted in blocking buffer for 1h at RT. Primary antibodies used in this study were: p-S129 aSyn (pSyn; CNDR mouse monoclonal IgG_2a_ 81A; 1:3,000); NeuN (Millipore A60; 1:3,000); MAP2 (CNDR rabbit monoclonal 17028: 1:3,000). This was followed by washing with PBS three times and incubating for 1h at RT in blocking buffer containing Alexa-Fluor-conjugated isotype-specific secondary antibodies (Thermo Fisher). Cells were washed three times with PBS and DAPI (0.4 μg/ml in PBS) was added to visualize cell nuclei. Stained cells were imaged using an InCell 2200 (GE Healthcare Life Sciences). Images from 9 fields were collected from each well with a 20x objective. Image analysis and quantification was performed using InCell Toolbox Analyzer software (GE Healthcare Life Sciences).

#### Statistical analysis

Data are shown as mean ± standard deviation unless stated otherwise. Statistical analysis for FLT and FRET experiments were conducted by an unpaired Student’s t-test using GraphPad to determine statistical significance for all experiments. Values of p-value < 0.05 were considered statistically significant.

Outcome measures for PFF induced neuron pathology (DxA 81A/NeuN ratio) were compared between experimental conditions using linear mixed-effects models, with fixed effects for condition and random effects for experiment, well by condition, and experiment by well by condition, to account for within-well correlation, variation by experiment, and variation of conditions across experiments. Outcome measures were log-transformed for analysis for approximate normality and homogeneity of variance across conditions and experiments. Estimated differences between conditions, along with their 95% confidence intervals, were exponentiated to report as ratios of the outcome measures between conditions. Analyses were conducted using R^*112*^ version 3.6.1 including the packages lme4^*113*^ version 1.1-21 and lmerTest^*114*^ version 3.1-1.

## Acknowledgements

We thank Nagamani Vunnam from the Sachs Group, Tory Schaaf, Samantha Yuen, Andrew Thompson, and Razvan Cornea from the Thomas Group and Benjamin Grant from Fluorescence Innovations, for technical support and discussions. Compound dispensing was performed at the UMN Institute of Therapeutic Drug Discovery and Development (ITDD) High-Throughput Screening Laboratory, and spectroscopy at the UMN Biophysical Technology Center. This research uses technology patented by the University of Minnesota, with an exclusive commercial license to Photonic Pharma LLC. The authors disclosed receipt of the following financial support for the research, authorship, and/or publication of this article: This study was supported by U.S. National Institutes of Health (NIH) grants to J.N.S. (NINDS grant 3R01NS084998_ and to J.N.S. and D.D.T. (NIA SBIR grant 1R43AG063675). This research was also supported by the NIH National Center for Advancing Translational Sciences, grant UL1TR002494. The content is solely the responsibility of the authors and does not necessarily represent the official views of the National Institutes of Health’s National Center for Advancing Translational Sciences.

## Author contributions

A.R.B. designed and conducted the experiments. E.E.L provided assistance with cell-based assays and western blot experiments. M.C.Y produce and purified recombinant protein. M.H. and K.L. performed primary neuron PFF assays. D.D.T. provided expertise on FRET and HTS, and provided comments and edits to the manuscript. M.E. and R.B. performed statistical model development and analysis on the PFF pathology model. E.E.L. provided comments and edits to the manuscript. A.R.B. and J.N.S. wrote the manuscript.

## Materials & Correspondence

Correspondence and material requests should be sent to Jonathan Sachs.

---

**Supplemental Fig. 1.**
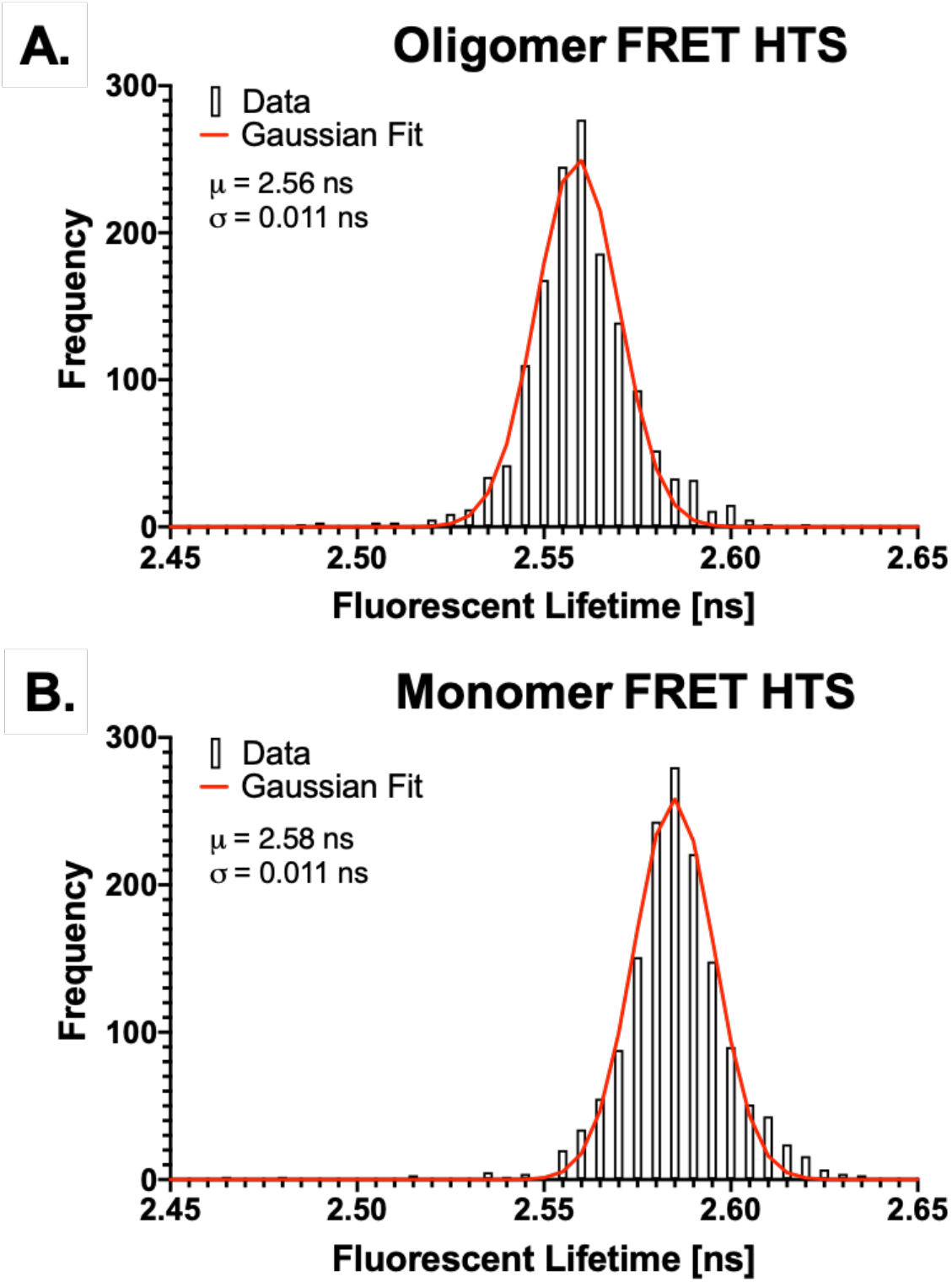
Gaussian Fit of Pilot HTS presented in Fig. 4. **(A)** Inter-molecular (oligomeric FRET HTS) and **(B)** intra-molecular (monomeric FRET HTS) pilot HTS both conform to good agreement with a gaussian FLT response. Mean and standard deviations were used to determine probable hit compounds that are prioritized for subsequent dose response FRET experiments

**Supplemental Fig. 2.**
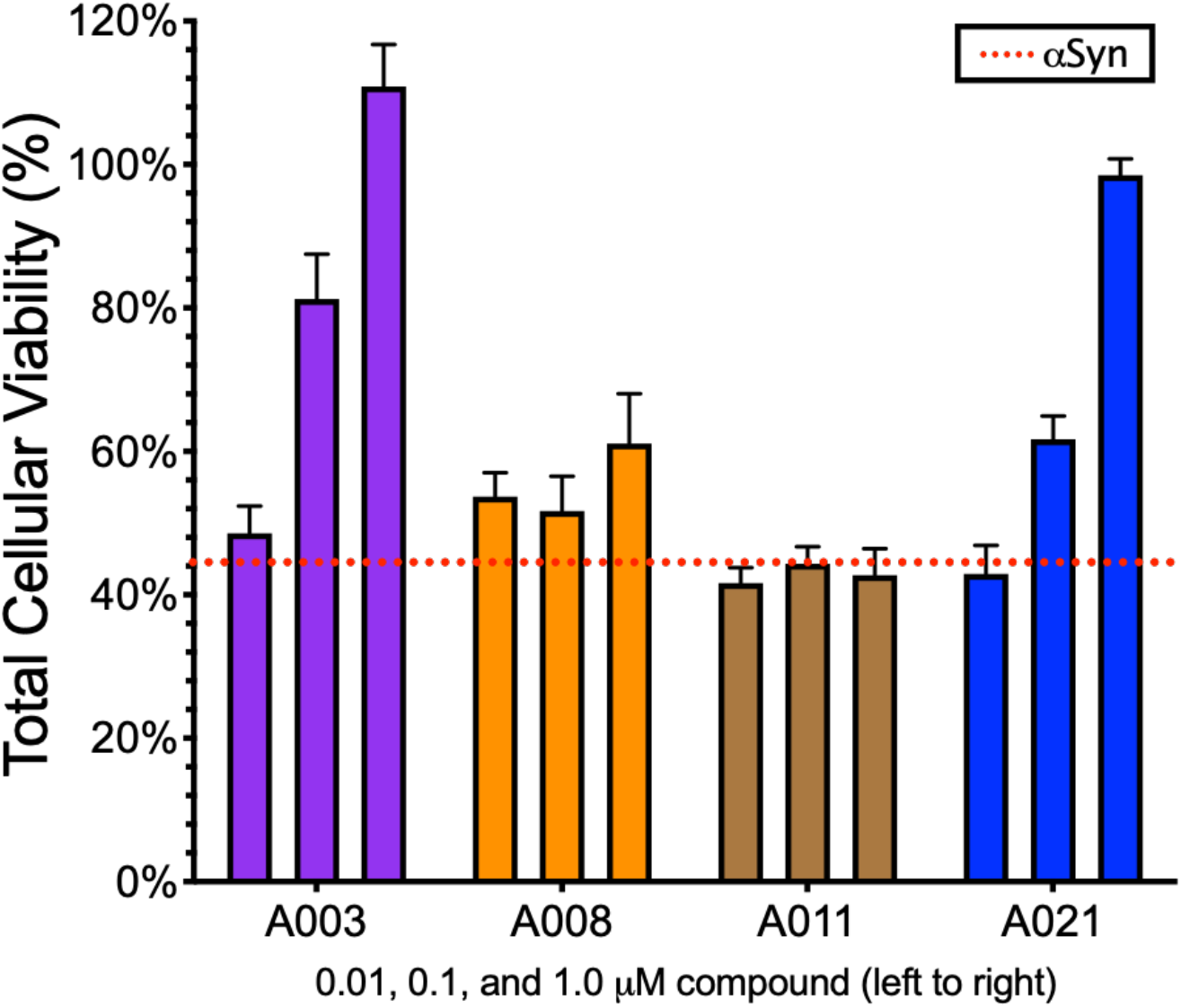
Rescue of aSyn induce cytotoxicity in SH-SY5Y cells. Overexpression of aSyn in SH-SY5Y cells results in reduced cellular viability (~44%, red dashed line) as determined by CytoTox-Glo cytotoxicity assay. A three-dose assay was used as a preliminary screen for hit-compounds. Not all FRET hits resulted in rescue of toxicity (e.g. A003 and A021 show strong response whereas A008 has only mild recovery and A011 show no rescue of toxicity.

**Supplemental Fig. 3.**
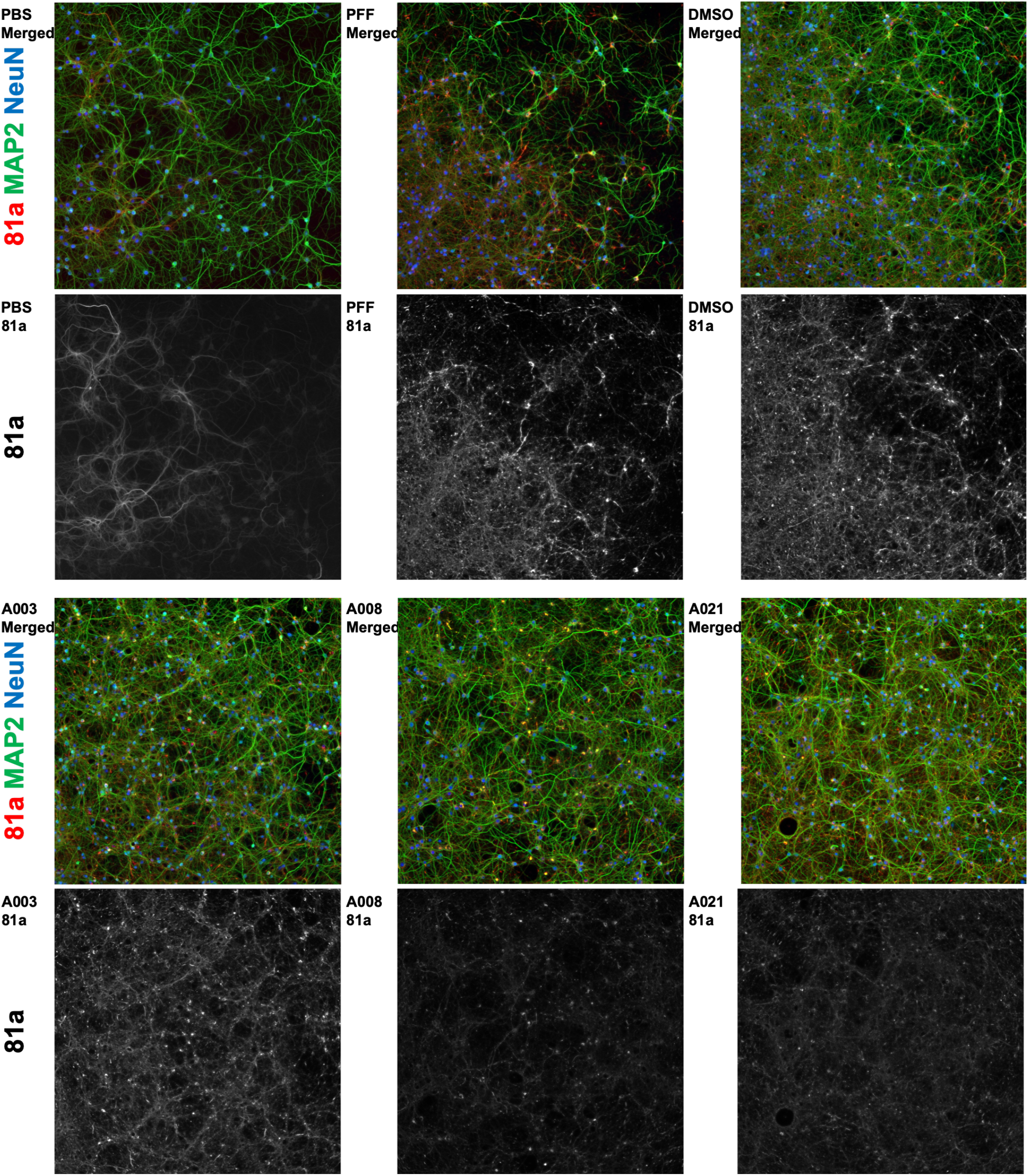
Rescue of aSyn PFF induced pathology in a primary neuron model. Using a msPFF aSyn induced pathology model in primary cortical mouse neurons we monitored phospho-S129 aSyn (81a) normalized by the neuron count (NeuN) as a readout for pathology. Neurons were treated with vehicle (PBS), PFF, PFF+DMSO, and three hit compounds (A003, A008, A021 @ 1uM) showed significant increase in 81a/NeuN signal whereas two hit compounds (A008 and A021) reduced the overall pathology load. Experiments were done in triplicate with three independent wells per experiment. Neurons were cultured for 7DIV, transfected with msPFF +/- drug, incubated 7DIV then processed for IFC. Quantification of 81a/NeuN signal is reported in **Fig. 6B.**

**Supplemental Fig. 4.**
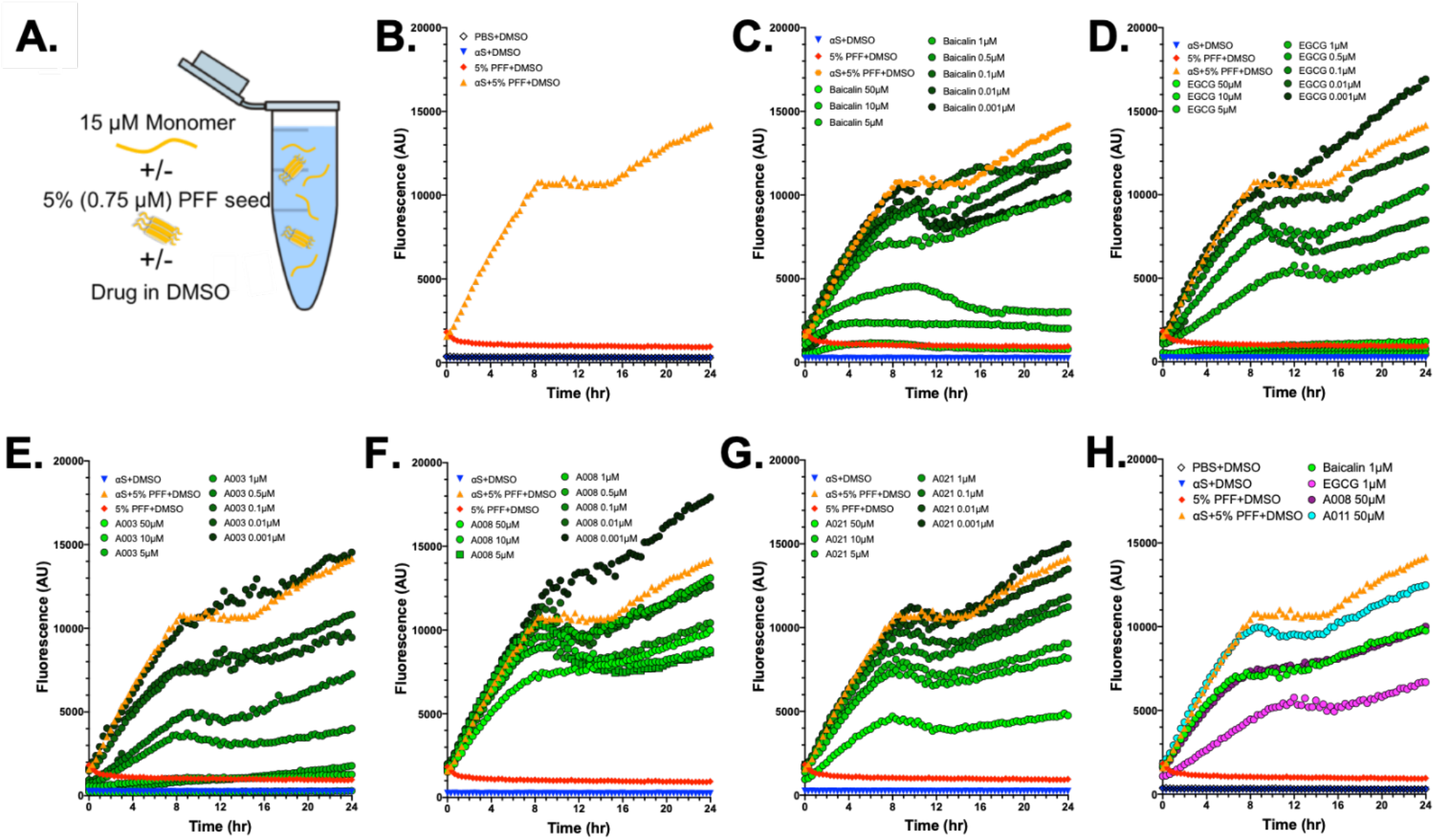
Seeded Thioflavin-T aggregation assay show direct protein interaction for subset of hit compounds. **(A)** Schematic of experiment. Mixtures of 15μM monomer aSyn with 5% (0.75μM) PFF were incubated with 20μM ThT and a range of hit compound concentrations. **(B)** Control samples are plotted in all graphs. aSyn monomer only (blue), 5% PFF (red), seeded monomer (orange) and PBS only Z(black) provide reference traces. **(C and D)** ThT CRC response for control compounds EGCG **(C)** and Baicalin **(D)**. Panels **(E-G)** are ThT CRC for A003, A008 and A021 respectively. Each of these traces were normalized to determine IC_50_ values for inhibition of seeded aSyn aggregation (see **Fig. 7C**). **(H)** Other compounds tested in ThT assay, A0011 show no aggregation inhibition. In panels B-H three control samples are included for reference: aSyn monomer only (blue); 5% aSyn PFF (red), and monomer + seed (yellow).

